# The evolution of non-motif selectivity determinants in *Monosiga brevicollis* PDZ domains

**DOI:** 10.1101/2020.05.28.121053

**Authors:** Melody Gao, Iain G. P. Mackley, Samaneh Mesbahi-Vasey, Haley A. Bamonte, Sarah A. Struyvenberg, Louisa Landolt, Nick J. Pederson, Lucy I. Williams, Christopher D. Bahl, Lionel Brooks, Jeanine F. Amacher

## Abstract

The evolution of signaling pathways is complex and well-studied. In particular, the emergence of animal multicellularity had a major impact on protein-protein interactions and signaling systems in eukaryotic cells. However, choanoflagellates, our closest non-metazoan ancestor, contain a number of closely related signaling and trafficking proteins and domains. In addition, because choanoflagellates can adopt a rosette-/multicellular-like state, a lot can be gained by comparing proteins involved in choanoflagellate and human signaling pathways. Here, we look at how selectivity determinants evolved in the PDZ domain. There are over 250 PDZ domains in the human proteome, which are involved in critical protein-protein interactions that result in large multimolecular complexes, e.g., in the postsynaptic density of neuronal synapses. Binding of C-terminal sequences by PDZ domains is often transient and recognition typically involves 6 residues or less, with only 2 residues specifying the binding motif. We solved high resolution crystal structures of *Monosiga brevicollis* PDZ domains homologous to human Dlg1 PDZ2, Dlg1 PDZ3, GIPC, and SHANK1 PDZ domains to investigate if the non-motif preferences are conserved, despite hundreds of millions of years of evolution. We also calculated binding affinities for GIPC, SHANK1, and SNX27 PDZ domains from *M. brevicollis*. Overall, we found that peptide selectivity is conserved between these two disparate organisms, with one exception, mbDLG-3. In addition, we identify 178 PDZ domains in the *M. brevicollis* proteome, including 11 new sequences, which we verified using Rosetta and homology modeling. Overall, our results provide novel insight into signaling pathways in the choanoflagellate organism.

## Introduction

Cellular processes that are a result of internal or external signals require an extensive network of intracellular communication. The transducers of these information pathways are proteins, whose specific three-dimensional structures are specialized for specific functions. Often, multiple protein domains exist in a single polypeptide chain, molecular “beads on a string” that together dictate the activities and binding partners of the protein in the cell (1). A particular pattern of this modular architecture can be exquisitely conserved in evolution for a specific protein family; for example, the SH3-SH2-Kinase Src tyrosine kinase architecture is a functionally relevant module that exists in uni- and multicellular species, whose last common ancestors existed almost one billion years ago (2, 3). Critically, each individual signaling domain, here, SH3, SH2, and kinase, is also used in a variety of different protein scaffolds (4). Thus, knowing the structure-function relationship for individual domains is also important in building a holistic understanding of the signaling networks of an organism.

An example of a signaling domain that is well conserved in evolution is the PDZ domain, named for the first proteins identified that contain a shared “GLGF” sequence, <u>P</u>SD-95, <u>D</u>lg1, and <u>Z</u>O-1 (5–9). The human proteome contains 272 PDZ domains in a variety of protein architectures, but all share the same general function. PDZ domains are approximately 80-100 residues, bind the extreme C-terminus of target proteins, and share a structural fold, consisting of a core antiparallel β-sheet and 1-2 α-helices (**Fig. 1**) (10–16). These are scaffolding domains that bind target proteins in order to facilitate localization to enzymatic auxiliary domains located on the same protein, or to mediate trafficking and signaling pathways, also determined by other protein-protein interactions at other domains in the protein or as a part of a larger macromolecular complex (11, 17, 18). An example of the scaffolding nature of PDZ domains is the postsynaptic density of neurons, where multiple receptor signaling networks are brought into close physical proximity due to a number of PDZ domain-mediated interactions (19).

**Figure 1.**
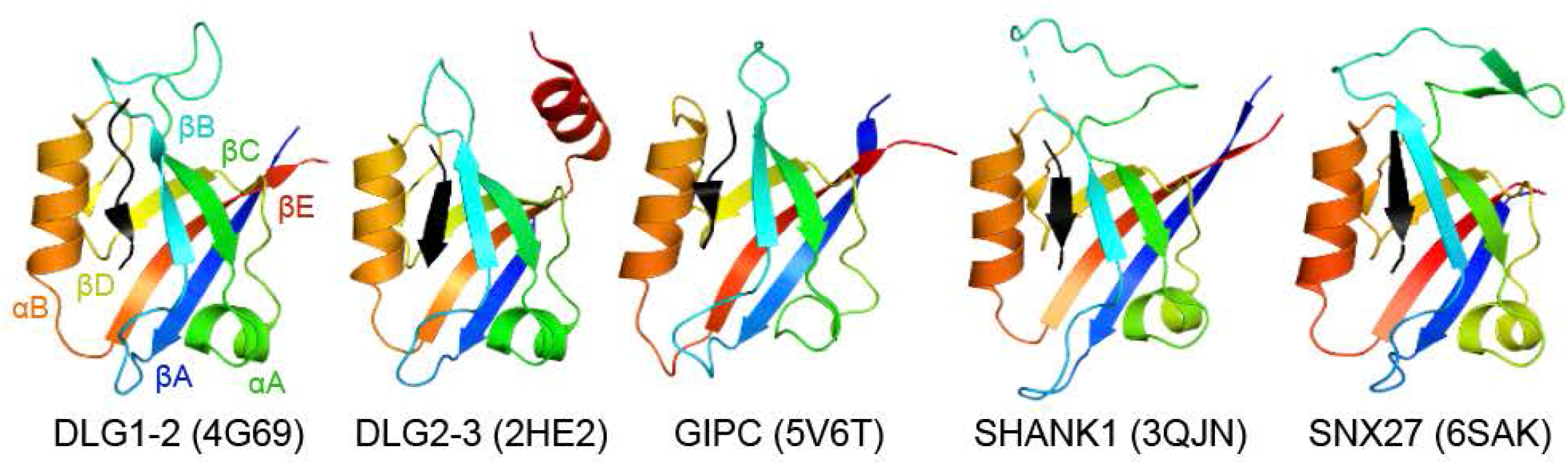
Conserved fold of PDZ domain structures. The human PDZ homologues of the *M. brevicollis* PDZ domains studied in this paper are shown in cartoon representation, colored by conserved secondary structure elements, as labeled. Bound peptides are in black cartoon.

Despite the importance of PDZ domains in cellular processes, teasing apart selectivity in this domain family can be a challenge. These domains bind to short sequences in target proteins, often interacting with only 6 amino acid residues. In fact, the motifs of classically determined PDZ binding classes are dependent on only two residues, the extreme C-terminal residue, termed P^0^ and two residues adjacent, or P^−2^ (11). Work in the last 10+ years using high throughput techniques, e.g., phage display, peptide array, or the *hold-up assay*, has shifted this classical view of PDZ domain binding to appreciate the importance of binding interactions at non-motif residues in the peptide-binding cleft (20–22). In addition, a number of elegant studies using directed evolution or other techniques have successfully altered PDZ selectivity, often through only a small number of amino acid substitutions or post-translational modifications (23–27).

Considering how relatively straightforward it is to vary PDZ target specificity and the prevalence of these domains throughout the tree of life, including in bacteria, yeast, and plants, it is perhaps not surprising that the evolution of specificity can also be traced on a case-by-case basis (26, 28, 29). However, what remains to be determined is whether or not the non-motif selectivity determinants in PDZ domains related by evolution are also conserved, despite vastly different levels of complexity in signaling pathways, e.g., in uni-versus multicellular organisms, or those with and without a nervous system. Previous work looking at the evolution of PDZ domains found that six amino acid positions determine lineage relationships amongst 40,000 PDZ domains in 40 proteomes and that four of these positions are in direct contact with non-motif peptide residues (P^−1^ and P^−3^) (30). This result suggests that homologous proteins will share positive and negative preferences for all residues that interact with the peptide-binding cleft, including those beyond the P^0^ and P^−2^ motif positions.

In order to investigate these questions on a molecular level, we crystallized and solved the structures of four PDZ domains from the choanoflagellate, *Monosiga brevicollis*, including homologues of PDZ domains from the human proteins Dlg1, GIPC1, and SHANK1 (**Fig. 1**). We also investigated the binding affinities of a homologue of human SNX27 (**Fig. 1**). These proteins are important in postsynaptic signaling and well conserved in *M. brevicollis*, despite over 200 million years of evolution between the last common ancestor of humans and choanoflagellates and the emergence of neurons (19, 31–33). Overall, choanoflagellates, such as *M. brevicollis*, are a powerful tool in studying cellular differentiation, considering they can switch between a uni- and multicellular-like *rosette* state (34–38).

Structural and binding affinity analyses confirm that non-motif selectivity is mostly conserved in these proteins, with a notable exception. In the third PDZ domain of the *M. brevicollis* Dlg1 homologue, which we refer to as mbDLG-3, there are substitutions in binding cleft residues that interact with non-motif residues, as well as a histidine-to-tyrosine substitution in a motif-determining position. This mutation is not shared in Dlg proteins from a number of other organisms, although it is in the Dlg protein from another choanoflagellate species, *Salpingoeca rosetta*, suggesting it is unique to choanoflagellates. Previous studies investigated the molecular basis of evolution, expansion, and *rewiring* in PDZ domain networks; however, here we find that for closely related PDZ domains, selectivity determinants for all residues in the binding cleft are generally conserved in evolution, despite a lack of conservation in shared target proteins (39, 40). In addition, we analyze the *M. brevicollis* proteome by conducting pairwise sequence alignments with all human PDZ domains in order to characterize its PDZome, verifying novel domains using homology modeling and Rosetta.

## Results

### Case Study I: Structural and biochemical characterization of mbGIPC PDZ

To determine if residues that directly interact with the ligand are conserved, including those that recognize non-motif positions in Class I PDZ domains, we set out to characterize a number of PDZ domains from *Monosiga brevicollis* with clear homology to human PDZ domain. We first chose to investigate the homolog of the human GAIP interacting protein, C terminus, or GIPC (41). GIPC was first identified as an interactor of the G-alpha interacting protein (GAIP), but was quickly shown to also interact directly with G-protein coupled receptors, as well as dopamine and NMDA receptors in excitatory synapses of the central nervous system (33, 41–43). Thus, GIPC is important for both G-protein coupled receptor (GPCR) and neuronal signaling in human cells, and additional studies have shown that it broadly regulates vesicular trafficking of many transmembrane receptors via interactions with myosin VI (14).

The presence of a GIPC homolog in *M. brevicollis* is consistent with the identification of adhesion GPCRs in choanoflagellates (44, 45). Overall, full length GIPC proteins from human and *M. brevicollis* (UniProt ID: A9VCZ3_MONBE, termed mbGIPC) share 56% sequence identity over 79% of the protein. The human GIPC protein is a Class I PDZ binding domain, defined as recognizing the general sequence motif X-S/T-X-ϕ(where X=any amino acid and ϕ=any hydrophobic amino acid) (11). Recognition of the P^−2^ Ser/Thr residue is facilitated by hydrogen bond formation with a conserved histidine in the first position of the conserved αB helix, termed αB-1 (11). Consistent with this, the human and choanoflagellate GIPC PDZ domains are 58% identical over 88% of the sequence, including shared carboxylate-binding loop sequences of ALGL and conservation of the Class I-defining histidine in the αB-1 position.

We expressed and purified mbGIPC PDZ using previously described methods, and as described in more detail in the Experimental Procedures and Supporting Information (22, 46). Briefly, we used recombinant expression in *Escherichia coli* cells, followed by affinity and size exclusion chromatography to produce purified mbGIPC PDZ protein. With protein in hand, we calculated binding affinities for human GIPC targets using fluorescence polarization, including GAIP, tyrosinase-related protein 1 (TYRP1), and the β-1 adrenergic receptor (B1AR) (**Table 1**). Experimental protocols were based on previously described methods, and are described in more detail in the Experimental Procedures and Supporting Information (22, 46–48).

**Table 1.** Binding affinities of mbGIPC PDZ.

|  |  | K<sub>i</sub> (μM) |
| --- | --- | --- |
|  | Sequence | mbGIPC |
| GAIP | QGPSQSSSEA | 0.140 ± 0.014 |
| TYRP1 | KLQNPNQSVV | 1.7 ± 1.0 |
| B1AR | RPGFASESKV | 2.6 ± 0.34 |
| mbGIPC C-term | VEPKKSFDEI | >1000 |

The binding affinities of mbGIPC PDZ for human GIPC PDZ targets suggest a large degree of conservation in selectivity determinants. Specifically, the affinity of mbGIPC PDZ for GAIP (C-terminal sequence: QSSSEA) is 140 nM, despite a BLASTP search revealing no obvious GAIP homolog in *M. brevicollis* (49). The binding affinities for TYRP1 (sequence: PNQSVV) and B1AR (ASESKV) are ~10 and 20x worse, respectively. These values are still relatively high as compared to typical PDZ domain interactions, which can range from the nanomolar to hundreds of micromolar range, but are centered around 1-30 μM (11, 48, 50). Neither TYRP1 nor B1AR have clear homologues in *M. brevicollis*, according to BLASTP (49). Thus, we determined the crystal structure of mbGIPC PDZ in order to directly compare the peptide-binding clefts of the mbGIPC and human GIPC PDZ domains.

We crystallized and solved the structure of mbGIPC PDZ to a high resolution of 1.2 Å, as described in the Experimental Procedures and Supplementary Information. Overall, this structure is consistent with the conserved PDZ fold, characterized by the central five-stranded antiparallel β-sheet (βA-E). As mentioned above, while many PDZ domains contains two α-helices (αA-B), it appears that αA is slightly strained and therefore not fully formed in the mbGIPC structure, although this is also true of the human GIPC PDZ domain (PDB IDs: 5V6B and 5V6T) (14). The peptide ligand forms an additional strand of the central β-sheet (**Fig. 2A**). Data collection and refinement statistics are in **Table S1A**.

**Figure 2.**
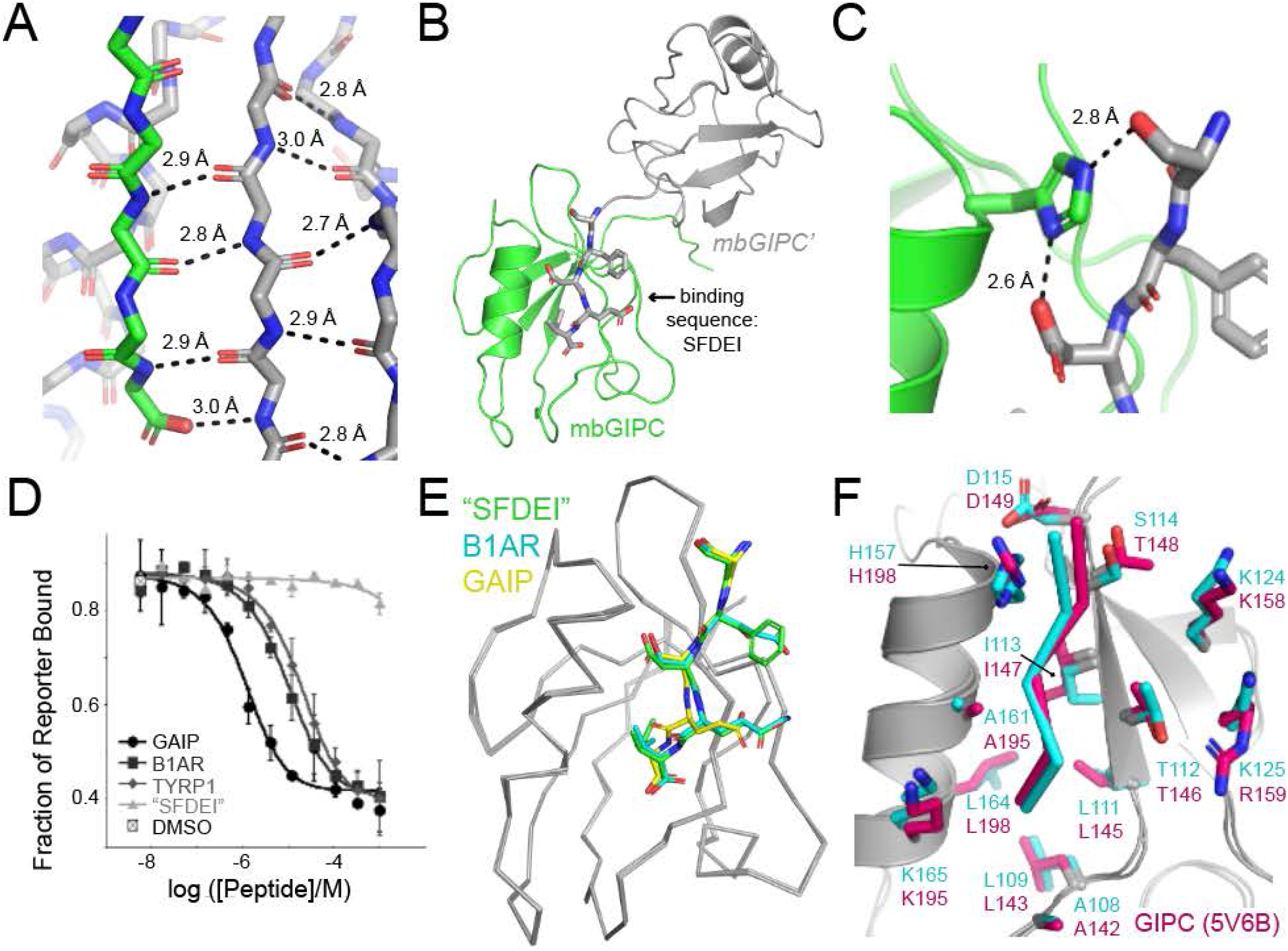
The crystal structure of the mbGIPC PDZ domain. (A) The interaction of mbGIPC (gray) with the C-terminal tail of a molecule related by symmetry (green), shown in stick, reveals a canonical PDZ-peptide interaction where the peptide forms an additional strand of an antiparallel β-sheet. Distances are labeled. (B) The spatial relationship between mbGIPC (green cartoon) PDZ and a molecule related by symmetry (gray cartoon, *mbGIPC’*) is shown. The last 5 residues of the *mbGIPC’* C-terminal tail (sequence SFDEI) are shown in gray stick. (C) The binding of mbGIPC (green cartoon, His157 side chain in stick representation) to the C-terminal tail of *mbGIPC’* (gray sticks) is unconventional and a crystal artifact. (D) Average fluorescence polarization displacement isotherms are shown for mbGIPC PDZ. Titration curves are shown for the following peptides: GAIP (circles), B1AR (squares), TYRP1 (diamonds), and a decameric peptide matching the C-terminal residues of the construct, ending in “SFDEI” (triangles). The average value from 6 DMSO controls is also shown (square with X). Error bars indicate standard deviation from the mean for duplicate experiments. (E) Alignment of mbGIPC PDZ domains with three separate C-terminal tail sequences (gray ribbon, RMSD = ≤ 0.21 Å for ~350 main chain atoms), with tail sequences as sticks and colored as labeled. (F) The conservation between mbGIPC (gray cartoon, with cyan side chain residues as sticks; peptide is in cyan ribbon) and human GIPC (PDB ID: 5V6B, with peptide from 5V6T (hot pink ribbon); gray cartoon with hot pink side chain residues as sticks). Residues in the peptide-binding cleft are labeled. All stick representation is colored by heteroatom (O=red, N=blue).

Although we had added the high resolution GAIP peptide during crystallization, we were surprised to see that our crystal structure lacked the bound peptide. Instead mbGIPC was interacting with the C-terminal tail of a molecule related by symmetry (**Fig. 2B**). This is a common mode of co-crystallization for PDZ domains and ligands, e.g., in the NHERF1 PDZ bound to the cystic fibrosis transmembrane conductance regulator (CFTR) structure (50). The NHERF1 PDZ1-CFTR example is distinct from our structure, however, in that the C-terminus of mbGIPC is not a Class I PDZ-satisfying motif (sequence: KSFDEI). In our structure, which we will refer to as mbGIPC_SFDEI_, we see that the P^0^ Ile is accommodated by a hydrophobic pocket, as expected in Class I PDZ interactions. However, the conserved αB-1 His residue is forming hydrogen bonds with the Asp in the P^−2^ position (distance: 2.6 Å), as well as the Ser in the P^−4^ position (2.8 Å) (**Fig. 2C**). Solution experiments using fluorescence polarization revealed that this interaction is a crystal artifact, mbGIPC_SFDEI_ does not bind a peptide mimicking the final 10 residues of mbGIPC_SFDEI_ (sequence: VEPKKSFDEI) with affinities < 1 mM (**Figs. 2D and S1A, Table 1**). A truncated mutant (mbGIPC_trunc_, lacking the final 7 residues of our original construct) reveals the same result (data not shown).

In order to investigate the stereochemistry of a peptide binding interaction with mbGIPC that is not an artifact of crystallization, we mutated the final 5 residues of our original construct to those matching B1AR (mbGIPC_B1AR_; C-terminal sequence: SESKV), GAIP (mbGIPC_GAIP_; SSSEA), and TYRP1 (mbGIPC_TYRP1_; NQSVV) (**Table S1A**). Previous work from ourselves and others suggests that the P^−5^ position is an important selectivity determinant in some PDZ domains (21, 22). However, we chose to keep this residue a lysine in our new constructs, due to apparent crystal lattice contacts with its side chain, suggesting it may be important for crystallization (**Fig. S1B**).

All 3 complexes successfully crystallized in the same space group as mbGIPC_SFDEI_ and we determined crystal structures of mbGIPC_B1AR_ and mbGIPC_GAIP_. The overall conformations of these structures to each other, as well as to the mbGIPC_SFDEI_ structure, were very similar, with pairwise structural alignment RMSD values ≤ 0.21 Å for ~350 main chain atoms (**Fig. 2E**). We were unable to fully refine the mbGIPC_TYRP1_ structure despite a successful molecular replacement solution, due to anisotropic data and relatively low resolution, compared to the others. Partial refinement (R_work_/R_free_ = 24.3/28.9) shows clear peptide-specific density, confirming that this sequence interacts with mbGIPC in a manner that is consistent with PDZ domain peptide binding (**Fig. S1C**). However, our structural analyses of the mbGIPC and human GIPC PDZ domains will be limited to the mbGIPC_B1AR_ and mbGIPC_GAIP_ structures.

Our mbGIPC structures share high structural similarity with the human GIPC PDZ domain. Structural alignment of main chain atoms between the mbGIPC_B1AR_ and hGIPC PDZ domains is 0.607 Å over 299 main chain atoms. This human GIPC PDZ structure was crystallized with the intracellular region of Plexin-D1 (C-terminal sequence: CYSEA) and the structures confirm that the peptide binding cleft of mbGIPC and human GIPC is very well conserved (**Fig. 2F**). We were unable to purify soluble human GIPC PDZ in our lab, but our data strongly suggests that the binding affinities would be similar between these domains, including preferences for amino acids at non-motif positions.

### Case Study II: Structure Characterization of mbSHANK1 PDZ

We previously compared binding affinities for another *M. brevicollis* PDZ domain, that of mbSHANK1 (UniProt ID: A9V7E4_MONBE), a protein that is homologous to human SHANK1 (46). In this work, we also created a homology model of mbSHANK1 PDZ using SwissModel and predicted stereochemical differences in the peptide binding pockets between these two proteins, specifically in those residues that interact with the P^−3^ position (46). Here, we expand that investigation by presenting the crystal structure of mbSHANK1 PDZ (**Fig. 3, Table S1B**).

**Figure 3.**
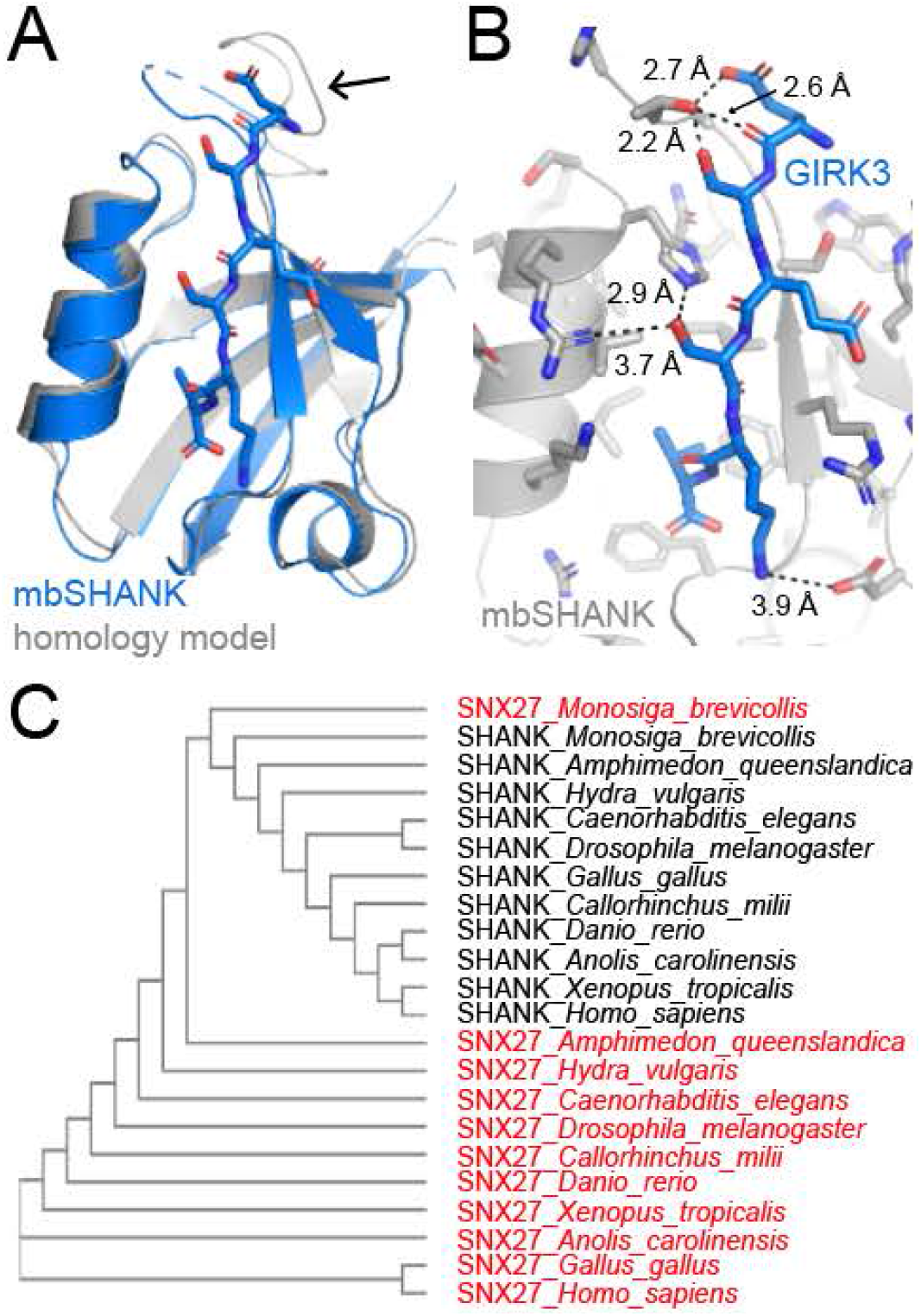
The crystal structure of the mbSHANK1 PDZ domain. (A) The mbSHANK1 structure (blue cartoon, peptide in stick representation) is very similar to a previously reported homology model (gray cartoon), RMSD = 0.668 Å over 276 main chain atoms (46). The black arrow highlights differences in the flexible βB-βC loop. All sticks are colored by heteroatom (O=red, N=blue). (B) The interactions of the mbSHANK1 PDZ domain (gray cartoon, with side chains as sticks) with the F*-GIRK3 peptide (blue sticks) is characteristic of PDZ-peptide interactions. Measurements between interacting residues in the peptide-binding cleft are labeled. (C) Phylogenetic tree showing the relationship of a number of SHANK and SNX27 PDZ domain sequences from 11 organisms. SNX27 sequences are colored red. The mbSHANK1 and mbSNX27 sequences sit at the branch point of the other SHANK and SNX27 sequences.

The protein mbSHANK1 PDZ was expressed and purified as previously described (46). Crystallization of this protein in complex with a fluoresceinated peptide matching the C-terminus of GIRK3 (F*-GIRK3, sequence: LPPPESESKV) is described in the Experimental Procedures and Supporting Information. We collected data to a high resolution of ~2.2 Å, however phasing by molecular replacement and structure refinement proved challenging. We employed an iterative Rosetta modeling approach coupled with Phenix in order to determine a molecular replacement solution with high confidence using Phaser (LLG = 1498, TFZ = 27.5), as described in detail in the Supplementary Information (51–54). Ultimately, our refinement difficulties were due to a large degree of anisotropy in the diffraction data. Specifically, the high-resolution limit along the a* and b* directions (2.2 Å) was substantially higher than that along the c* direction (3.4 Å). We were ultimately able to refine this model by truncating and scaling the reflections file appropriately, using the UCLA-DOE Diffraction Anisotropy Server (55). Notably, crystallization attempts with a fluorescent β-PIX peptide (sequence: NDPAWDETNL), despite binding mbSHANK1 PDZ with 62-fold higher affinity, were unsuccessful (**Table 2**). We were also unable to grow crystals of mbSHANK1 in the apo form or following incubation with non-fluorescent versions of either the β-PIX or GIRK3 peptides.

**Table 2.**
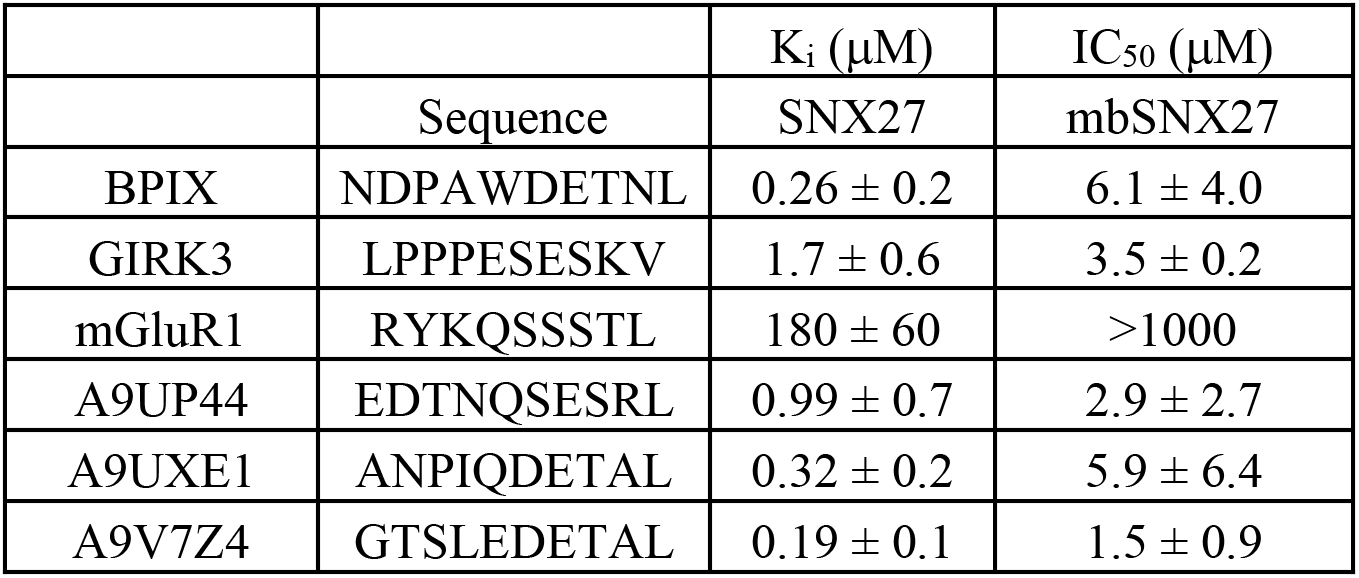
Binding affinities of SNX27 and mbSNX27 PDZ domains.

|  |  | K<sub>i</sub> (μM) | IC<sub>50</sub> (μM) |
| --- | --- | --- | --- |
|  | Sequence | SNX27 | mbSNX27 |
| BPIX | NDPAWDETNL | 0.26 ± 0.2 | 6.1 ± 4.0 |
| GIRK3 | LPPPESESKV | 1.7 ± 0.6 | 3.5 ± 0.2 |
| mGluR1 | RYKQSSSTL | 180 ± 60 | >1000 |
| A9UP44 | EDTNQSESRL | 0.99 ± 0.7 | 2.9 ± 2.7 |
| A9UXE1 | ANPIQDETAL | 0.32 ± 0.2 | 5.9 ± 6.4 |
| A9V7Z4 | GTSLEDETAL | 0.19 ± 0.1 | 1.5 ± 0.9 |

The crystal structure of mbSHANK1 bound to F*- GIRK3 is structurally very similar to our previously determined homology model (46). The overall RMSD of these two structures is 0.668 Å over 276 main chain atoms, with the largest discrepancies occurring in the flexible βB-βC loop (**Fig. 3A**). In our structure, we see non-covalent interactions between T471 and the side chains of the P^−4^ Ser and P^−5^ Glu residues, as well as the P^−5^ Glu carbonyl, which may have helped to stabilize the βB-βC loop for crystallization, although six residues of the loop are disordered in our structure (**Fig. 3B**). In addition, we see electrostatic interactions between D488 and the P^−1^ Lys, as well as H517 and R518 with the P^−2^ Ser (**Fig. 3B**).

In our previous work and based on our mbSHANK1 homology model, we hypothesized that the modest increase in affinity for β-PIX by mbSHANK1 PDZ (K_i_ = 13 μM versus 20 μM for human SHANK1 PDZ) was due to an additional arginine residue near the P^−3^ position (46). Our experimental structure, however, reveals that neither of the arginine residues in the vicinity are interacting with the P^−3^ Glu of GIRK3. It is unclear from our mbSHANK1:GIRK3 structure what may account for this slight difference in affinity, however the crystal structure confirms that the peptide-binding cleft is conserved as we previously reported, with the exception of the previously-described residues near the P^−3^ peptide position (46). In addition, the sequence and length of the βB-βC loop, which directly interacts with peptides in our structure, varies quite dramatically: The N-terminal residues of the 11-residue loop for mbSHANK1 PDZ are THSAAEQT, while for human SHANK1 PDZ, an 18-residue loop, they are KAQTPIEE.

When we ran our initial BLASTP search for SHANK1 PDZ homologues in *M. brevicollis*, the top two sequence hits were relatively close in sequence identity: A9V7E4_MONBE, with 34% sequence identity over 93 residues, as well as A9URU5_MONBE, with 36% sequence identity over 89 residues. Sequence alignments using the full-length A9V7E4_MONBE protein searching the human proteome confirmed its homology to the SHANK protein family, specifically due to the additional presence of ankyrin repeat domains, as well as SH3 and SAM domains (31, 56). Sequence alignments using the full-length A9URU5_MONBE sequence and human proteome suggested that it is a homologue of sorting nexin-27 (SNX27), with 25% sequence identity over 96% of the protein. Therefore, we will refer to A9URU5_MONBE as mbSNX27.

We were still interested in the relationship between the PDZ domain sequences in these four proteins, and conducted a phylogenetic tree analysis of 10 PDZ sequences for SHANK1 or SNX27 homologues in a variety of organisms, as well as the PDZ domain sequences of mbSHANK1 and mbSNX27 (**Figs. 3C**). Because we see that mbSHANK1 and mbSNX27 sit at the branch point between the SNX27 and SHANK1 sequences, we expressed and purified SNX27 and mbSNX27 PDZ domains, as described in the Experimental Procedures and Supporting Information, and compared binding affinities for all four domains using fluorescence polarization to 6 decameric peptides matching the C-termini of: β-PIX, GIRK3, and mGluR1, as well as A9UP44_MONBE, A9UXE1_MONBE, and A9V724_MONBE, which were previously identified as potential *M. brevicollis* targets of mbSHANK1 (46) (**Fig. S2A-C, Table 2**).

Our results reveal that overall, peptides which bind human SNX27 PDZ with relatively high affinity also bind mbSNX27 PDZ very strongly. We measure similar results with human SHANK1 and mbSHANK1 PDZ domains (46). However, in all cases, the exact order of highest to lowest affinity peptides is distinct, perhaps reflective of single substitutions in the peptide binding cleft. We described the differences for SHANK1 and mbSHANK1 PDZ domains above and previously (46). A homology model of mbSNX27, using SNX27 PDB as a template (PDB ID: 6SAK) contains the following substitutions at residues that may interact with the peptide (numbering based on SNX27): R58K, V61T, A83H, and R122I (**Fig. S2D**). In addition, while the mGluR1 peptide binds the human PDZ domains with moderate affinity, it shows no measurable affinity for either of the *M. brevicollis* PDZ domains. Taken together, the resulting binding affinities are consistent with our hypothesis that mbSHANK1/human SHANK1 and mbSNX27/human SNX27 each share common ancestors.

### Case Study III: Structural characterization of PDZ domains in mbDLG

We also wanted to investigate multiple PDZ domains from a single *M. brevicollis* protein. Therefore, we experimentally determined the structures of PDZ2 and PDZ3 from UniProt ID A9UT73_MONBE (**Table S1B**). This protein is most closely related to the discs large family of human proteins, sequence identity values for the full-length protein are as follows for: human Dlg1 (35.5%), Dlg2 (35.0%), Dlg3 (34.1%), and Dlg4 (30.5%). Therefore, we will refer to these PDZ domains as mbDlg-2 and mbDlg-3, respectively. The highest domain-to-domain identity for both mbDLG-2 and mbDLG-3 is to PDZ2 and PDZ3 of human Dlg2 (**Table S2**).

Expression, purification, and crystallization of mbDLG-2 and mbDLG-3 were performed as described in the Experimental Procedures and Supplementary Information, and following standard protocols. Interestingly, the sequence of mbDLG-2 contains zero aromatic residues, and thus has an extinction coefficient at λ=280 nm of 0. Therefore, protein quantification for crystallization was done via SDS-PAGE analysis using bovine serum albumin as a standard, as described in the Experimental Procedures and Supporting Information (**Fig. S3A**). The mbDLG-2 was incubated with a peptide matching the final 10 residues of the HPV16 E6 oncoprotein (sequence: SSRTRRETQL) prior to crystallization and crystallized in two distinct space groups: *P* 2_1_ 2_1_ 2_1_ and *I* 2 (which is related to *C* 2) (57). A structure of human Dlg2 PDZ2 (PDB ID: 2BYG) was successfully used as a search model in molecular replacement for both crystal forms.

The mbDLG-2 structure is very similar in both crystal symmetries. The overall RMSD is 0.153 Å for 300 main chain atoms (**Fig. 4A**). We are able to resolve an additional 6 C-terminal residues in the orthorhombic (*P* 2_1_ 2_1_ 2_1_) crystal, which form an α-helix, and there is positive density in the peptide-binding cleft that likely corresponds to the HPV16 E6 peptide (**Fig. 4B**). Iterative rounds of refinement after placing peptide residues into this density confirm that there may be multiple confirmations of the peptide within the pocket and that its occupancy is likely 0.50 or less. Therefore, we were not confident in modeling peptide residues. In the centered monoclinic (*I* 2) crystal, we are able to resolve the entire βA-βB loop, however, crystal contacts with molecules related by symmetry are not compatible with peptide binding (**Fig. 4C**). The major difference between these two structures is the location of the carboxylate-binding loop, a shift that is consistent with carboxylate-binding loop flexibility in a number of apo and peptide-bound structures of human Dlg2 PDZ2 (**Fig. 4A, S3B**). Comparison with the peptide binding-cleft of human Dlg2 PDZ2 (PDB ID: 4G69) reveals only two relatively conservative differences: N339S and K392R (**Fig. 4D**).

**Figure 4.**
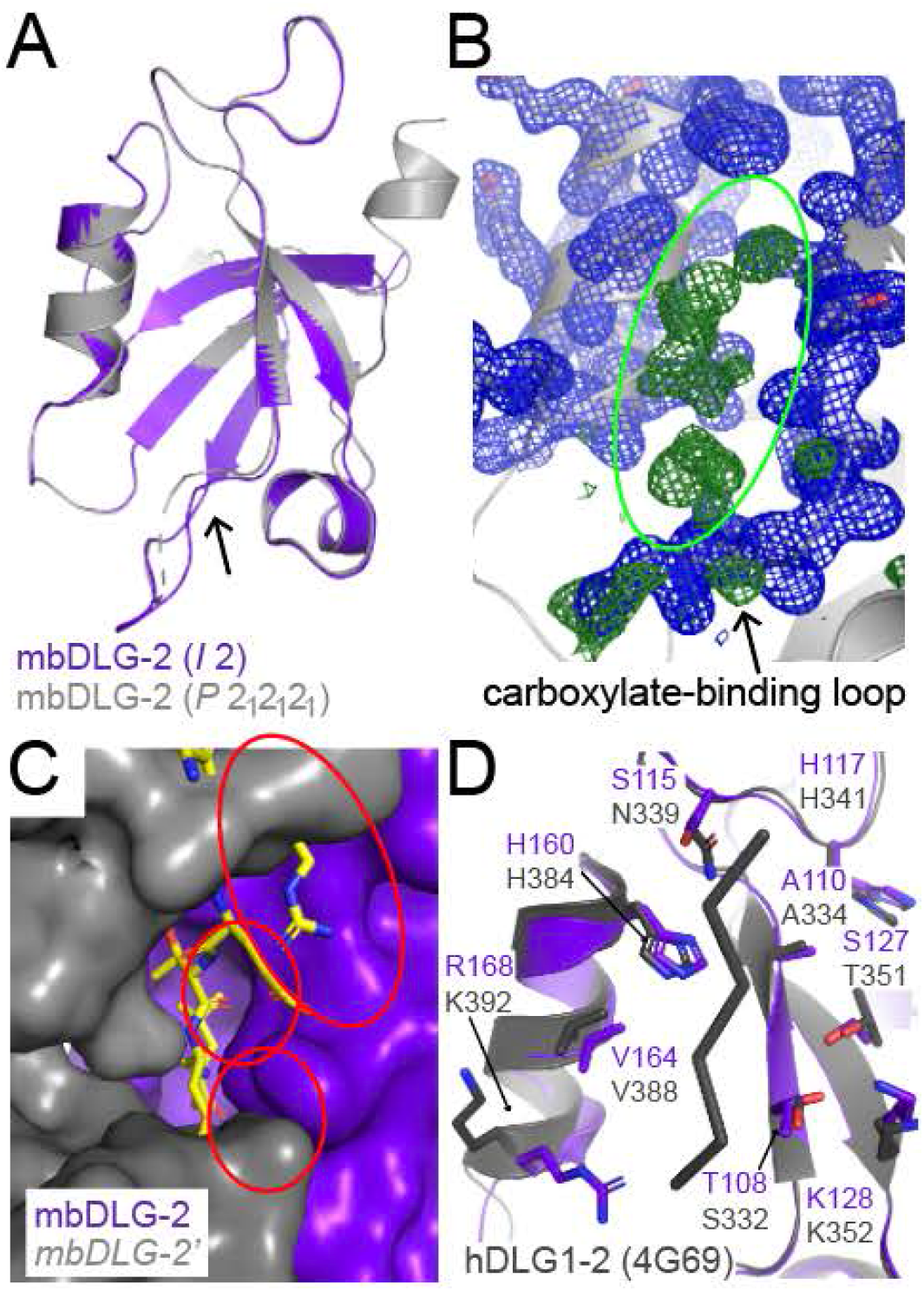
The crystal structure of the mbDLG-2 PDZ domain. (A) Alignment of the two mbDLG-2 structures, which crystallized in different space groups: *I* 2 (purple cartoon) and *P* 2_1_ 2_1_ 2_1_ (gray cartoon). Overall RMSD = 0.153 Å for 300 main chain atoms. The biggest difference between the structures is a shift in the carboxylate-binding loop, indicated by a black arrow. (B) The *P* 2_1_ 2_1_ 2_1_ mbDLG-2 PDZ domain (gray cartoon, electron density is shown in blue mesh, 2*F*_o_-*F*_c_ map contoured at 1σ) structure revealed strong positive density (green mesh, *F*_o_-*F*_c_ map contoured at 2.5σ, highlighted by the green circle) in the peptide binding cleft, which is likely the HPV16 E6 peptide that was incubated with the PDZ domain prior to crystallization; however, iterative rounds of refinement suggested that the occupancies of peptide atoms were < 0.5 and that there were multiple conformations. Ultimately, the peptide could not be confidently modeled. A black arrow points to the carboxylate-binding loop, which is labeled. (C) The mbDLG-2 PDZ domain (purple surface) that crystallized in the *I* 2 space group is not bound by peptide, due to molecules related by symmetry (gray surface). The mbDLG-2 PDZ domain was aligned with the CAL PDZ domain structure bound to a HPV16 E6 peptide (PDB ID: 4JOP, RMSD = 0.859 Å for 239 main chain atoms). Steric clashes between molecules related by symmetry and the HPV16 E6 peptide (yellow sticks) are highlighted with red circles. (D) The conservation between mbDLG-2 (purple cartoon, with side chain residues as sticks) and human Dlg1 PDZ2 (PDB ID: 4G69, gray cartoon with side chain residues as sticks and peptide as ribbon). Residues in the peptide-binding cleft are labeled. All stick representation is colored by heteroatom (O=red, N=blue).

Structure determination of the mbDLG-3 PDZ domain was less straightforward. A number of structures were used as molecular replacement search models, including multiple human Dlg1, Dlg2, and Dlg3 PDZ3 structures, as well as models determined using *de novo* structure prediction by the Robetta server, which uses the Rosetta fragment insertion method (58). Ultimately, we were able to determine a borderline solution with PSD-95 (or Dlg4) PDZ3 (PDB ID: 6QJF), LLG = 57 and TFZ = 7.2. This model was improved using Phenix Autobuild, which allowed us to generate a suitable model for further refinement (59). Clear electron density for unmodeled residues confirmed the lack of phase bias and the structure was refined to a final *R*_work_/*R*_free_ = 15.8/16.9 (**Table S1B**). Structural alignments with the 5 lowest energy models from Robetta revealed a range of RMSD values for main chain atoms between 0.902 Å (234 atoms) and 1.373 Å (259 atoms) (**Fig. S4A**). In contrast, the RMSD value for the alignment of mbDLG-3 and the A-protomer of 6QJF (PSD-95 PDZ 3) was 0.726 Å over 225 main chain atoms (**Fig. S4B**). Notably, although the βB-βC loop is twice as long in mbDLG-3 (16 residues versus 8 residues in PSD95 PDZ3), the N- and C-terminal residues of this loop follow a similar trajectory as 6QJF, which is not the case in any of the Robetta models (**arrows in Fig. S4A-B**).

Sequence alignments of mbDLG-3 with the human Dlg1-4 PDZ3 domains suggest that the Class I-determining αB-1 histidine residue is a tyrosine in mbDLG-3. Alignment of mbDLG-3 to a peptide-bound Dlg2 PDZ3 (PDB ID: 2HE2) structure, with an RMSD = 0.779 Å over 225 main chain atoms, confirms this class-switching difference (**Fig. 5A**). Critically, the P^−2^ Thr residue in the bound ETSV peptide sterically clashes with the tyrosine hydroxyl in mbDLG-3 (**Fig. 5B**). In general, the peptide binding cleft of mbDLG-3 is the most dissimilar to that of its nearest homologs, Dlg2 and PSD-95/Dlg4 PDZ3, with substitutions at four residues (numbering is for Dlg2 PDZ3): N434S, V436I, F448R, and H480Y (**Fig. 5C**).

**Figure 5.**
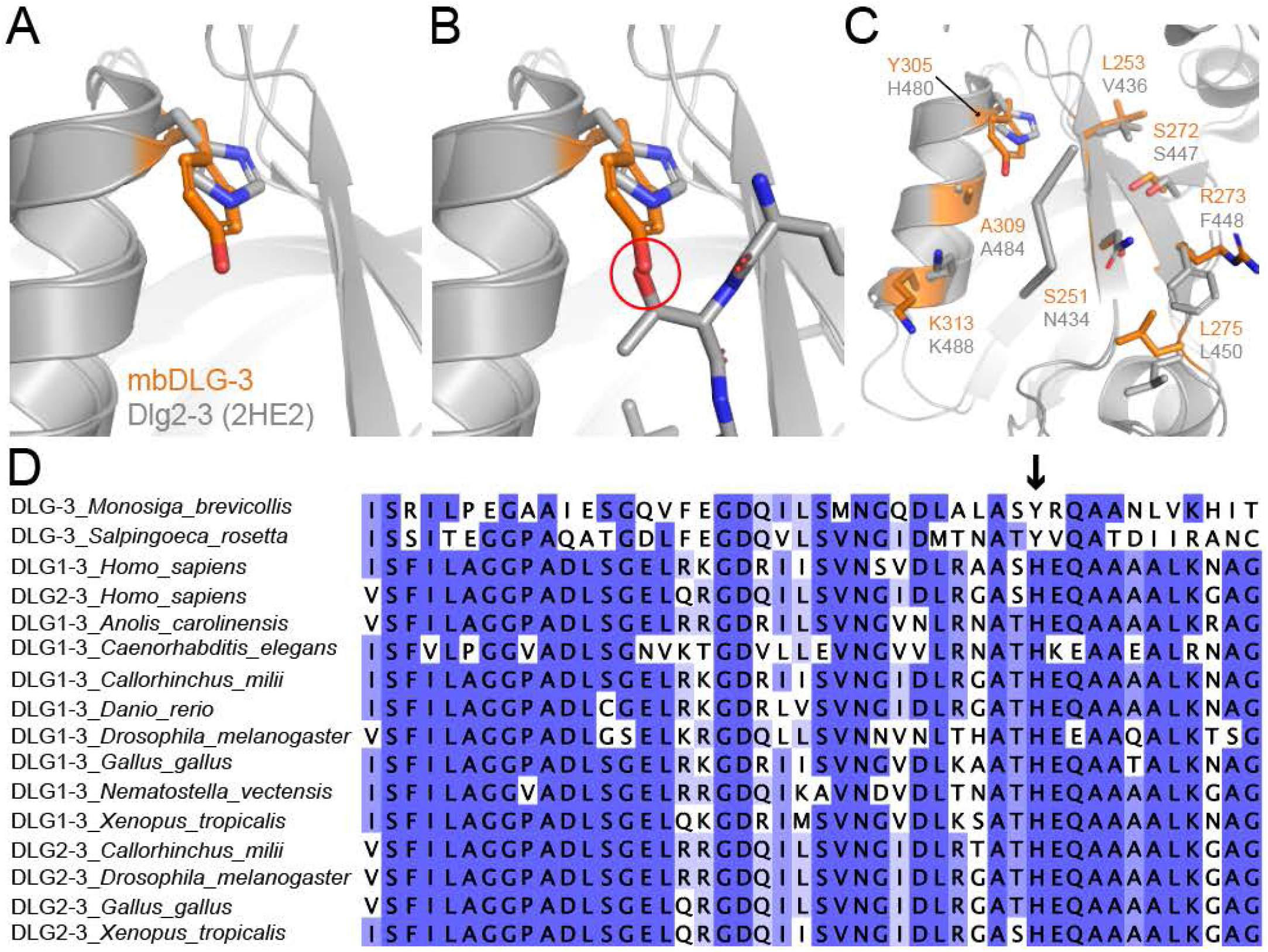
The crystal structure of the mbDLG-3 PDZ domain. (A-B) Substitution in the binding class-determining αB-1 residue between mbDLG-3 (gray cartoon, with the side chain of Y305 as orange sticks) and human Dlg2 PDZ3 (Dlg2-3, PDB ID: 2HE2; gray cartoon with the side chain of H480 as sticks). The P^−2^ Thr that interacts with H480 (gray sticks in (B)) in Dlg2-3 sterically clashes with Y305 in mbDLG-3 (red circle). (C) The conservation between mbDLG-3 (gray cartoon, with side chain residues as orange sticks) and human Dlg2-3 (gray cartoon with side chain residues as sticks and peptide as ribbon). Residues in the peptide-binding cleft are labeled. All stick representation is colored by heteroatom (O=red, N=blue). (D) Sequence alignment of Dlg1 PDZ3 and Dlg2 PDZ3 domains from multiple organisms reveals that the only PDZ domains with a tyrosine in the αB-1 position are those from choanoflagellate species, *Monosiga brevicollis* and *Salpingoeca rosetta*. This position is highlighted with a black arrow, and sequence alignment coloring is by overall percentage identity (darker blue = higher % identity).

We further investigated a number of Dlg sequences in other organisms to see if there are others with a H-to-Y substitution at the αB-1 residue. None of the Dlg1 or Dlg2 (where available) sequences from *Gallus gallus* (chicken), *Danio rerio* (zebrafish), *Callorhinchus milii* (shark), *Xenopus tropicalis* (frog), *Caenorhabditis elegans* (worm), *Anolis carolinensis* (lizard), or *Nematostella vectensis* (sea anemone) contain a tyrosine residue at the αB-1 position (**Fig. 5D, S4C**). In addition, sequence alignments with human PDZ domains that do contain an αB-1 Tyr all reveal significantly less sequence identity with mbDLG-3 than do the alignments to the human Dlg1-4 PDZ3 domains (44-49% over 62-78 residues); these include CYTIP (28% over 25 residues), DPTOR (25% over 75 residues), NHRF3-4 (29% over 75 residues), PDZIP-3 (29% over 75 residues), RADIL (34% over 77 residues), RHG21 (29% over 112 residues), and RHG23 (28% over 71 residues). Taken together, the αB-1 Tyr in mbDLG-3 appears to be a substitution unique to this Dlg protein in choanoflagellates, although the molecular basis for this change is not known. Future experiments to determine cellular targets of mbDLG-3 in *M. brevicollis* or *S. rosetta* would be interesting in order to decipher the functional consequence of this change.

### Identification of 178 PDZ domains in Monosiga brevicollis proteome

Finally, we wanted to determine the number of PDZ domains in the *Monosiga brevicollis* proteome. Previous work from 2010 identified 113 PDZ domains in 58 genes, using a combination of sequence alignment (i.e., HMMER 2.3.2) and evolutionary approaches (i.e., EvolMap); these results included human orthologs to well-studied human proteins such as Dlg, GIPC, and SHANK, the PDZ domains that are investigated extensively in our above case studies (30). However, more recent annotations from the UniProt database reveal 169 PDZ domains in 70 unique proteins (60, 61). Automatic annotation in UniProt is done using the EMBL InterPro system, as well as UniRule and the Statistical Automatic Annotation System (SAAS), which incorporate predictive models based on a number of sequence databases and machine learning (60–63).

For our search, we used a BLASTP-based approach, comparing all sequences in the *M. brevicollis* proteome to our 272 curated human PDZ domain sequences (11). We then wrote a Python-based program to filter the results by alignment length and to identify the top “hit” by sequence identity for each of our putative PDZ domains (**Table S3**). Our search also reported the alignment start and end residues, so that we could easily identify multiple PDZ domains in a single protein. We restricted ourselves to looking at results with alignment lengths of 65 residues or higher (excluding gaps), in order to reduce false positives. Manual curation of our results confirmed that this was a good cut-off because we successfully identified all of the annotated UniProt PDZ domains, as well as a number of additional ones. Using homology and Rosetta modeling we were able to further investigate our true and false positives, as described below. Overall, using our approach we identify a total of 180 PDZ domains in 77 proteins within the *M. brevicollis* proteome. In addition, we concluded that 2 PDZ domains annotated by UniProt are missing critical components of the conserved structural fold and are likely not PDZ domains, as described below. Our list therefore includes a total of 178 PDZ domains in *M. brevicollis*, including 11 previously unidentified PDZ domains.

### Rosetta and homology modeling of M. brevicollis PDZ domains

To validate our *M. brevicollis* PDZ domains, we first manually curated all sequence hits of alignment lengths of 65 or higher. The main criteria for this first step of filtering was the identification of a potential “GLGF” carboxylate-binding loop sequence in the vicinity of the N-terminal alignment residue. Next, we validated new and borderline *M. brevicollis* PDZ domains using Rosetta and homology modeling. “Borderline” sequence hits were classified as having a putative “GLGF” sequence, despite perhaps aligning with 4 or less of our 272 human PDZ domain sequences or having relatively high BLAST E-values, defined as >1E-5. This value is based on previously determined threshold cut-offs (49, 64). Notably, the top sequence identity reported for a given *M. brevicollis* PDZ domain may be from an alignment with a BLAST E-value >1E-5; in these cases, manual validation confirmed that a number of alignments with other PDZ domains are below the threshold (**Table S3**).

We used homology modeling with various PDZ templates and/or modeling using Rosetta to see if the conserved PDZ structural fold was consistent with each borderline sequence (65–70). All borderline sequences were verified using the homology modeling application in Rosetta (71). To improve sidechain packing, the predicted models were subjected to an all-atom refinement protocol using Rosetta (72–74). This approach identified a number of new candidate PDZ domains, defined as containing hydrophobic residues on the interior and hydrophilic residues on the exterior of the protein, as well as most of the conserved secondary structure elements, and positioning of the “GLGF” sequence in the correct location at the N-terminal end of the βB strand (11). We colored the structures by hydrophobicity of each residue, using *color_h.py* for PyMOL, and compared to the other choanoflagellate structures presented in this work (**Fig. 6**) (75). Interestingly, one of our PDZ hits is a protein with a carboxylate-binding loop sequence of “HWNL” (A9V1Y4); while an Asn residue in the 3^rd^ position of the “GLGF” loop would be a very rare occurrence, our previous analysis of human PDZ domains annotated in UniProt identified three human proteins, RAPGEF6, CAR11, and CAR14, with PDZ domains that have a Gln residue at this position (sequences: PLQF, TSQL, and LEQI, respectively), suggesting that an Asn can be accommodated (11).

**Figure 6.**
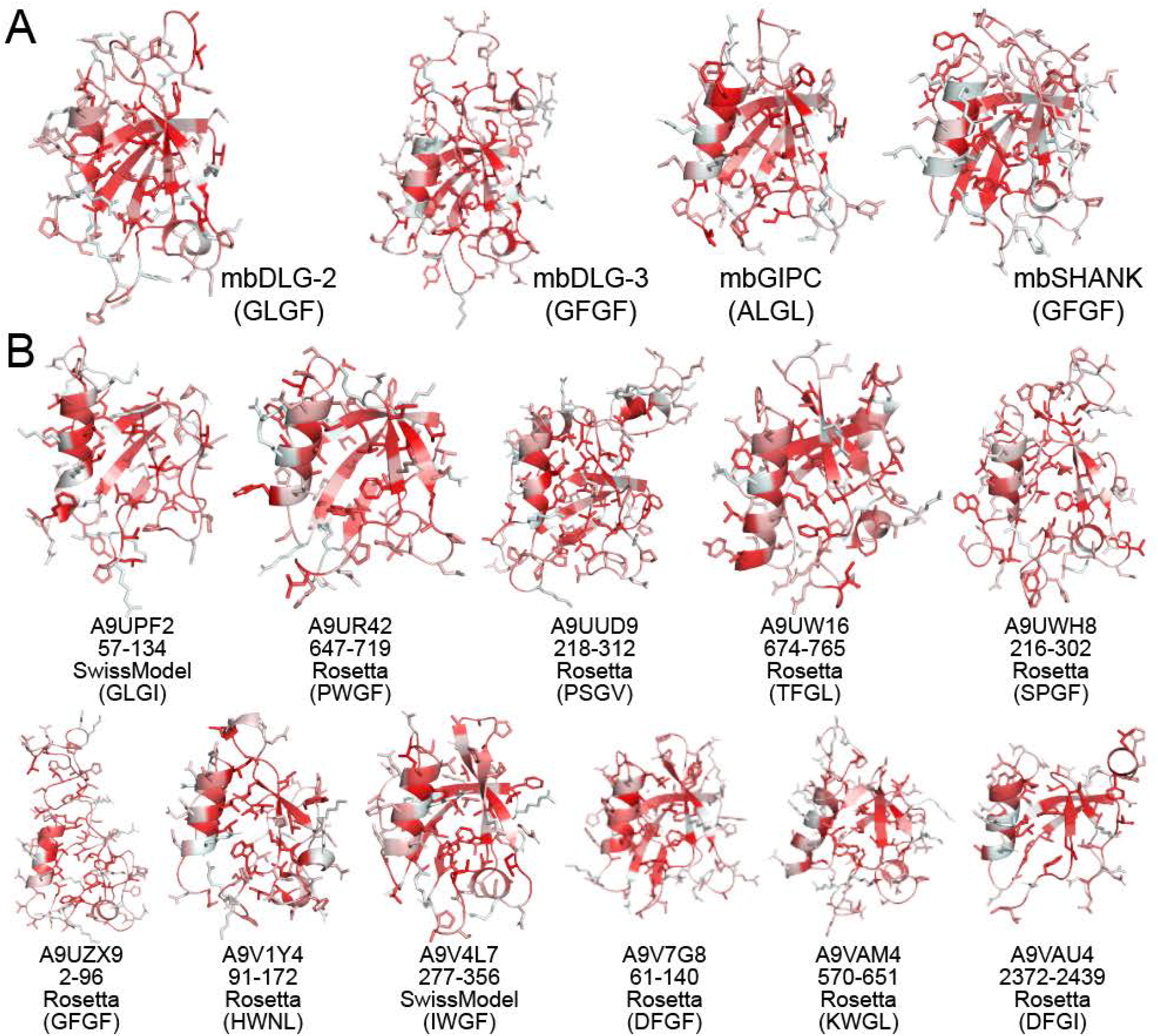
Hydrophobicity of choanoflagellate PDZ domains: experimentally-determined crystal structures (A) and models determined by Rosetta or SwissModel (B). All PDZ domains, with experimentally determined structures (A) or models of borderline sequences from our *M. brevicollis* proteome search (B) are labeled and shown in cartoon representation, with side chain sticks. Coloring is by degree of hydrophobicity, using previously determined values and *color_h.py* for PyMOL (75). The carboxylate-binding loop sequences are in parentheses under the protein name (A) or UniProt ID, residue numbers, and method for modeling (B). Overall, our models confirm that most hydrophobic residues are buried in the interior of the protein core and there are no buried hydrophilic residues, suggesting that the PDZ domain fold is reasonable for these sequences.

We also investigated 4 sequences that are annotated as PDZ domains by UniProt, but were flagged as borderline based on our criteria (**Fig. S6, Table S3**). The A9UPI8 PDZ18 Rosetta model contains two lysine residues near the interior of the protein, but these side chains may be in different orientations in an experimental structure (**Fig. S6A**). In addition, neither our SwissModel nor Rosetta models include the C-terminal residues of A9V109 PDZ1, after αB, and it is unclear if βE would properly form (**Fig. S6B**). Based on our analyses, A9UPI8 PDZ18 (residues 2222-2300) and A9V109 PDZ1 (residues 537-595) are likely PDZ domains, although further experiments would need to confirm these results. For the other two domains, the 3rd PDZ domain of A9V625 is defined as only 40 residues in UniProt (499-539), and our model suggests that while a longer sequence may adopt the PDZ fold, the carboxylate-binding loop sequence is “NQRC,” which is highly unusual (**Fig. S6C**). Therefore, we conclude that A9V625 PDZ3 is likely not a true PDZ domain. We also conclude that A9V7P9 does not contain a true PDZ domain. Here, our PDZ sequence alignment program only identifies a single alignment for this protein, even without length constraints considered: ZO1-2 at 31.6% identical over 76 residues (residues 409-482), with an E-value of 2.9 (**Table S3**). In addition, multiple attempts to determine homology or Rosetta models using residues ~380-470 or ~400-490 were unsuccessful, with a reasonable model reflecting a non-PDZ fold (**Fig. S6D**). Overall, our results suggest that a targeted BLASTP approach with a query consisting of all of the sequences from a human domain family, in combination with Rosetta and/or homology modeling, is a good approach for identifying protein domains in the proteomes of distantly related organisms.

## Discussion

Comparison of the selectivity determinants in specific peptide-binding domains and interactions in organisms related by hundreds of millions of years of evolution provides insight into signaling networks in those species. Here, we chose to use structural biology and biochemistry to investigate three PDZ domain-containing proteins that are important in human neuronal signaling in a species of choanoflagellates, our closest non-metazoan ancestors. Many of the human targets of SHANK1, GIPC1, and the Dlg family (including PSD-95) are either not conserved in choanoflagellates or do not contain PDZ binding sequences, as discussed in our previous work (46). Overall, we find that the motif and non-motif selectivity determinants are generally conserved in these related domains. Specifically, we see strong binding affinity correlations in SHANK1 and mbSHANK1 versus SNX27 and mbSNX27 PDZ domains, despite binding cleft substitutions in both cases. A notable exception is in mbDLG PDZ3, where we see a number of amino acid substitutions that directly affect target specificity. Presumably, this is due to unique signaling pathways in choanoflagellates, and future work should use computational or experimental methods to identify endogenous targets of mbDLG PDZ3, as well as the other studied PDZ domains (46, 76).

Our structures of four unique *M. brevicollis* PDZ domains provide the first structural determination of choanoflagellate PDZ domains to our knowledge. Furthermore, our comparisons with known human PDZ domain structures, as well as homology and Rosetta modeling confirm that because the PDZ domain fold is so well conserved, it is possible to get an initial idea of a PDZ domain structure without experimental structure determination. For example, we can use homology modeling with the closest-related human PDZ domains by sequence identity, to propose the structures of mbDLG-1 (template: INADL-8 (PDB ID: 2DM8), 49.3% identical over 69 residues) and mbSNX27 (template: SNX27 (PDB ID: 6SAK), 45.7% identical over 92 residues) (**Fig. 7A-B and S2, Table S3**). We can also propose the structures of PDZ domains that are in proteins with no obvious relation to human proteins, aside from the presence of one or more known domains, e.g., A9VDV9_MONBE, which contains 1 PDZ and 1 Kinase domain (according to UniProt). The highest sequence identity of this protein to any human protein, using BLASTP, is to the ROR2 tyrosine kinase receptor, at 26.53% over the kinase domain residues. The human ROR2 receptor does not contain a PDZ domain. As a template we used the PARD3-2 PDZ domain structure (PDB ID: 2KOM), which is 38.2% identical to the PDZ domain in A9VDV9_MONBE over 68 residues (**Fig. 7C**). We hypothesize that these types of analyses can be applied to PDZ domains from multiple organisms related by evolution.

**Figure 7.**
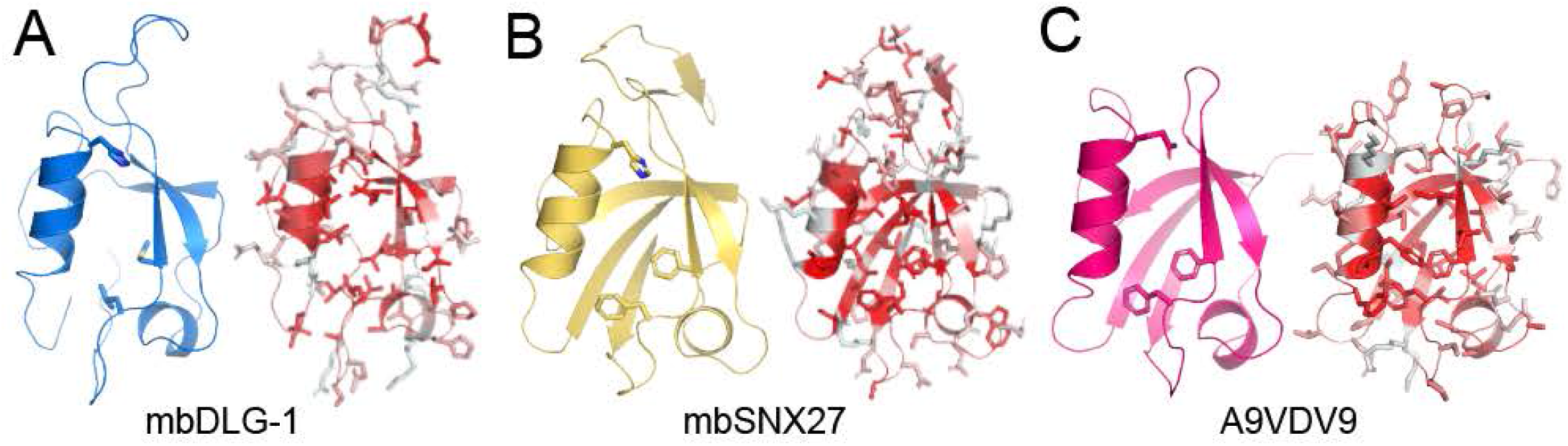
Homology modeling of *M. brevicollis* PDZ domains. Our results suggest that homology modeling is a reasonable tool for predicting the structure of PDZ domains in humans or other organisms, with no experimentally-determined structures. Here, we show homology models for (A) mbDLG-1 (template PDB ID: 2DM8), (B) mbSNX27 (template: 6SAK), and (C) A9VDV9_MONBE (template: 2KOM). For all, a cartoon representation of the model is on the left, with the αB-1 residue and carboxylate-binding loop side chains shown as sticks, and the hydrophobicity plot (cartoon with side chain sticks, residues colored by hydrophobicity) is on the right. The hydrophobicity plots confirm that the PDZ fold is reasonable for each of these sequences (i.e., hydrophobic residues in the protein core and hydrophilic/polar residues on the surface). Coloring for the left-side figures are: mbDLG-1 (blue), mbSNX27 (gold), and A9VDV9 (hot pink). All sticks are colored by heteroatom: O=red, N=blue, S=yellow.

Protein-protein interactions that involve PDZ domains act as critical nodes for signaling and trafficking pathways in a cell. It is clear that this is true in differentiated cells, such as those in complex multicellular organisms, as well as in single-celled organisms. Deciphering the PDZ-mediated interactions in choanoflagellates may elucidate important characteristics of the selectivity determinants and the evolution of this important peptide-binding domain. Furthermore, there are a number of proteins and protein architectures that contain PDZ domains in choanoflagellates that are not conserved in humans. Future work could investigate how these proteins, for example A9VDV9 mentioned above, act in signaling pathways in *M. brevicollis* and how this provides insight into the transition from uni- to multicellular life on Earth.

## Experimental Procedures

All protein expression and purification was performed as previously described (22, 46). Briefly, all PDZ domain coding sequences were inserted into the pET28a+ vector (GenScript) with an N-terminal His-tag and cleavable PreScission site. Additional details are in the Supporting Information. Crystallization experiments were done using hanging drop vapor diffusion and all conditions, which are described in the Supporting Information, originated from screening conditions in the Hampton Research PEG/Ion and PEG/Ion 2 screens. All crystal diffraction data was collected at the Advanced Light Source (ALS) at Lawrence Berkeley National Labs (LBNL). Fluorescence polarization experiments were performed as previously described (22, 47, 48, 77). Additional information on all experimental procedures is in the Supporting Information. PDB accession codes for the structures presented here are: 6X1X (mbGIPC_SFDEI_), 6X20 (mbGIPC_B1AR_), 6X22 (mbGIPC_GAIP_), 6X23 (mbSHANK), 6X1P (mbDLG-2, spacegroup *I* 2), 6X1N (mbDLG-2, spacegroup *P* 2_1_ 2_1_ 2_1_), and 6X1R (mbDLG-3).

## Acknowledgements

The authors would like to sincerely thank Drs. Christine Gee (UC Berkeley) and Dean Madden (Geisel School of Medicine at Dartmouth) for important discussions about anisotropic crystal data. In addition, Jacob Olson and Bodi Van Roy were involved in early work on mbDLG PDZ domain purification, and we would also like to thank the other members of the Amacher lab for helpful discussions and assistance. All data collection was done at the Advanced Light Source (ALS) at Lawrence Berkeley National Lab (LBNL) on beamline 5.0.1. We would like to thank the Berkeley Center for Structural Biology (BCSB) for assistance using the facility, specifically, Stacey Ortega for administrative and Marc Allaire and Dr. Daniil Prigozhin for technical help. The Berkeley Center for Structural Biology is supported in part by the Howard Hughes Medical Institute. The Advanced Light Source is a Department of Energy Office of Science User Facility under Contract No. DE-AC02-05CH11231. The Pilatus detector on 5.0.1. was funded under NIH grant S10OD021832. Other grant information: JFA was funded by NSF CHE-1904711, MG was funded by NSF-REU grant CHE-1757629, and CB was funded by Rosetta Licensing Fund grant RC8010. Start-up funds from Western Washington University also contributed to this project.

## Author contributions

JFA, CDB, and LB designed the experiments. MG, IM, SMV, HB, SS, LL, NP, and LW performed experiments. MG, IM, SMV, HB, SS, LL, NP, and LW contributed to data analysis. JFA wrote the manuscript and prepared the figures. All authors contributed to editing of the manuscript.

## Declaration of interests

The authors declare no competing interest.

